# The CUTIN SYNTHASE enzyme family was a key driver of cuticle emergence in land plants

**DOI:** 10.64898/2026.03.25.714142

**Authors:** Samuel Knosp, Florence Bernardeau, Elodie Lim, Giulia Zizi, Ludivine Malherbe, Mathieu Erhardt, Bénédicte Bakan, Hugues Renault

**Affiliations:** IBMP | Institut de biologie moléculaire des plantes, CNRS, University of Strasbourg, Strasbourg, France; INRAE, Biopolymers, Interactions, Assemblies Research Unit, La Géraudière, Nantes, France

**Keywords:** Adaptation, development, bryophytes, polymers, GDSL

## Abstract

The plant cuticle is a key adaptation acquired during the colonization of land. It forms a hydrophobic barrier at the interface with the environment, fulfilling essential protective and developmental functions. Despite its evolutionary significance and central role in land plant biology, the determinants that drove the emergence of the cuticle remain poorly understood. Here, we show that the CUTIN SYNTHASE (CUS) enzyme family, which synthesises the lipidic polyester that forms the structural framework of the cuticle in flowering plants, originated in a common ancestor of land plants, concomitant with terrestrialization. Using the moss *Physcomitrium patens*, we further demonstrate that CUS function is conserved among land plants. Inactivation of *CUS* genes disrupts gametophore development, the first tissue forming a cuticle during the moss life cycle, and compromises cuticle integrity. We also show that *P. patens* CUS enzymes localize to the apoplast, where they mediate the formation of a 10,16-dihydroxyhexadecanoic acid polyester using 2-mono-(10,16-dihydroxyhexadecanoyl)glycerol as substrate. Overall, our results reveal the conservation of CUS catalytic and physiological functions over 500 million years and support a pivotal role for this enzyme family in the emergence of the cuticle in an ancestral land plant during terrestrialization.

**SIGNIFICANCE:** The cuticle is a hallmark of land plants that fulfills essential roles, ranging from protection to development. Elucidating the mechanisms underlying its emergence and formation therefore has the potential to reveal fundamental aspects of land plant evolution and biology. Here, we show that the CUTIN SYNTHASE (CUS) enzyme family, which catalyzes the formation of the cuticle framework in flowering plants, arose in an ancestor of land plants during terrestrialization. Using the moss *Physcomitrium patens*, we further demonstrate that CUS function has been conserved for 500 million years across bryophytes and tracheophytes. We propose that the emergence of the CUS family was a key event in establishing the plant cuticle during terrestrialization.

## INTRODUCTION

Modern land plants (embryophytes) trace their origin to a singular evolutionary event termed terrestrialization, during which a lineage of streptophyte algae progressively adapted to life on land (Cheng et al. 2019; Carrillo-Carrasco et al. 2025). A key adaptation that facilitated this transition to desiccating terrestrial habitats was the specialization of the epidermal cell wall into a hydrophobic layer known as the cuticle (Renault et al. 2017; Kong et al. 2020; Philippe et al. 2020). Acting as an impermeable modification of the cell wall at the plant-environment interface, the cuticle establishes a diffusion barrier that limits the loss of water and solutes (Yeats and Rose 2013). Accordingly, the cuticle plays vital roles in plant resilience to abiotic constraints such as water deficit and high temperature, and also acts as a barrier to pathogen infection (Yeats and Rose 2013). Beyond these protective functions, increasing evidence points to important roles for the cuticle in plant development, ranging from preventing organ fusion during organogenesis to regulating anisotropic cell growth and maintaining hypoxic niches in meristems (Wellesen et al. 2001; Renault et al. 2017; Zhang et al. 2024; Raggi et al. 2026; Voloboeva et al. 2026).

The cuticle of flowering plants (angiosperms) is organized around cutin, a lipophilic polymer which is deposited at the outer face of the epidermal cell wall (Fich et al. 2016). This cutin matrix constitutes the structural framework of the cuticle and can be coated and impregnated with a range of hydrophobic compounds collectively referred to as waxes, which further reinforce its hydrophobic properties (Samuels et al. 2008). In tomato fruits and Arabidopsis flowers, the core component of cutin is a polyester of 10,16-dihydroxyhexadecanoic acid (10,16-diOH-C16:0) (**Fig. 1A**) (Isaacson et al. 2009; Li-Beisson et al. 2009). This polyester is synthesized by apoplastic enzymes known as CUTIN SYNTHASE (CUS), belonging to the GDSL lipase/esterase superfamily (Girard et al. 2012; Yeats et al. 2012). CUS enzymes catalyse the transesterification of 10,16-diOH-C16:0 at both terminal and midchain hydroxyl groups, using 2-mono-(10,16-dihydroxyhexadecanoyl)glycerol (2-MHG) as substrate (Yeats et al. 2012; Philippe et al. 2016, 2025), thereby generating a reticulated polymeric network (**Fig. 1A**). Each reaction cycle releases one molecule of glycerol, with a single glycerol moiety remaining attached to the initial monomer (**Fig. 1A**). Consistent with its central role in cuticle establishment, CUS deficiency has adverse effects on cuticle integrity and leads to detrimental tissue permeability in tomato fruit and Arabidopsis flowers (Girard et al. 2012; Yeats et al. 2012; Hong et al. 2017; Philippe et al. 2025). To date, CUS enzymes have not been investigated *in planta* beyond angiosperm models. Whether their key role in cuticle formation is conserved across land plants thus remains unknown.

**Figure 1.**
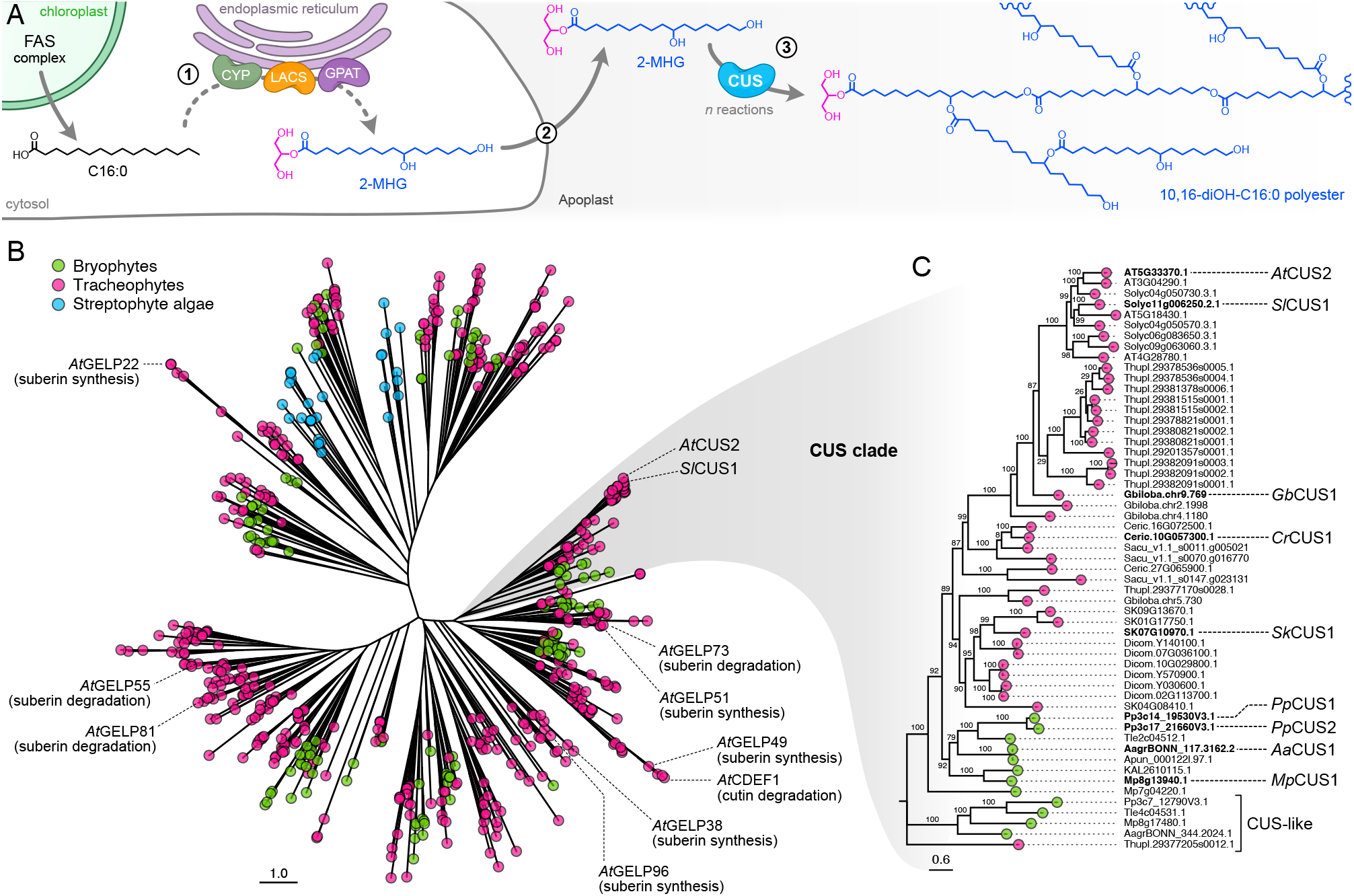
The CUTIN SYNTHASE enzyme family arose in embryophytes. (**A**) Schematic overview of the CUTIN SYNTHASE (CUS) metabolic pathway leading to polymerization of 10,16-dihydroxyhexadecanoic acid (10,16-diOH-C16:0) from 2-mono-(10,16-dihydroxyhexadecanoyl)glycerol (2-MHG) precursors in angiosperms. The pathway comprises three main steps: (1) biosynthesis of 2-MHG from hexadecanoic acid (C16:0) through the coordinated activity of ER-bound cytochromes P450 monooxygenases of the CYP86 and CYP77 families (Li-Beisson et al. 2009), long-chain acyl-CoA synthetase (LACS) (Schnurr et al. 2004; Bessire et al. 2007), and glycerol-3-phosphate acyltransferase (GPAT) (Li et al. 2007; Petit et al. 2016), (2) export of 2-MHG to the apoplast, at least partially mediated by ABCG transporters (Elejalde-Palmett et al. 2021), and (3) CUS-catalyzed transesterification of 10,16-diOH-C16:0 using 2-MHG as substrate. Esterification may occur at both terminal and midchain hydroxyl groups. Note that the precise sequence of enzymatic reactions in step 1 remains unresolved. (**B-C**) Reconstruction of the evolutionary history of the CUS enzyme family. (**B**) Maximum-likelihood protein tree (IQ-TREE3, Q.PFAM+R10) describing the phylogenetic relationships among 610 GDSL homologs derived from 19 streptophyte species. Proteins of relevance to the study are highlighted. CDEF1, Cuticle Destructing Factor 1; GELP, GDSL-type esterase/lipase protein. (**C**) Subtree highlighting the monophyletic CUS clade. Branch support values are based on 10,000 ultrafast bootstrap replicates. Scale bars indicate the number of amino acid substitutions per site.

In this study, we reconstructed the evolutionary history of the CUS enzyme family using recently available genomes spanning the diversity of Viridiplantae and show that this family arose in an ancestor of land plants, during the transition to terrestrial environments. We further demonstrate that CUS function in the moss *Physcomitrium patens*, from catalytic properties to physiological roles, is conserved. Altogether, our findings support a central role for the CUS enzyme family in the emergence of the cuticle in an ancestral embryophyte during terrestrialization.

## RESULTS

### The CUS family arose in an ancestor of embryophytes

An earlier study reported that the CUS family is present in tracheophytes and the bryophyte *Physcomitrium patens* (Yeats et al. 2014). However, the limited availability of plant genome data at that time, particularly from streptophyte algae, precluded the precise determination of the family’s origin. To address this, we surveyed for CUS homologs in the genomes of 23 Viridiplantae species, including two chlorophytes, seven streptophyte algae, six bryophytes and eight tracheophytes (**Supp. Fig. S1**). Using the characterized tomato *Sl*CUS1 protein as a bait (Girard et al. 2012; Yeats et al. 2012) and a phmmer E-value cut-off of e^-20^, we retrieved 610 protein homologs, all belonging to the GDSL superfamily. No homologs were detected in the two chlorophyte genomes or in the streptophyte algae *Mesostigma viride* and *Chlorokybus atmophyticus* (**Supp. Fig. S1**). We then reconstructed the evolutionary history of CUS based on the 610 retrieved sequences. The resulting phylogenetic tree showed that the two characterized CUS proteins from tomato and Arabidopsis (*Sl*CUS1 and *At*CUS2) (Girard et al. 2012; Yeats et al. 2012; Hong et al. 2017) were not closely related to streptophyte algal homologs (**Fig. 1B**). Instead, they fell within a monophyletic clade of embryophyte proteins containing homologs from all examined tracheophyte and bryophyte genomes (**Fig. 1C**). This indicates that the CUS family originated in an ancestor of embryophytes and has been strictly conserved across this lineage. Four bryophyte and gymnosperm homologs were positioned at the base of the CUS clade (**Fig. 1C**). They were tentatively annotated as CUS-like proteins as these sequences were more loosely related to characterized CUS homologs and were not consistently present across the surveyed species. The phylogenetic reconstruction also identified a CUS sister clade containing proteins involved in both cutin and suberin metabolism (GELP73, GELP51, GELP49, CDEF1; **Fig. 1B**) (Takahashi et al. 2010; Ursache et al. 2021; Philippe et al. 2025), suggesting shared GDSL-mediated biosynthetic mechanisms between these two lipid polymers. Other GELPs previously implicated in suberin synthesis or degradation (Ursache et al. 2021) were scattered throughout the phylogenetic tree (**Fig. 1B**), highlighting the complexity of the GDSL superfamily.

### *CUS* inactivation alters *P. patens* gametophore development

To date, no *in planta* studies of *CUS* genes have been conducted in models other than angiosperms. We therefore investigated the two *CUS* paralogs from the moss *P. patens* identified in the phylogenetic analysis (Pp*CUS1* and Pp*CUS2*; **Fig. 1C**) to assess functional conservation within the family. Bulk RNA-seq data showed that both genes are expressed at very low levels in protonema (**Fig. 2A**). Prominent expression appeared in gametophores, the first vegetative tissue forming a cuticle during the moss life cycle, with Pp*CUS1* being the most expressed paralog (**Fig. 2A**). At later developmental stages, Pp*CUS* genes were primarily expressed in archegonia and young sporophytes (**Fig. 2A**). To further refine spatial expression patterns, we generated knock-in promoter-reporter lines for both genes (**Supp. Fig. S2**). GUS assays showed that Pp*CUS1* and Pp*CUS2* promoters had overlapping expression domains, with strongest activity in gametophore apices and the basal regions of phyllids (**Fig. 2B**). Extended GUS incubation, however, detected promoter activity throughout the entire gametophore (**Supp. Fig. S2**). Next, we investigated the consequences of *CUS* inactivation by isolating single and double CRISPR/Cas knock-out lines (**Supp. Fig. S3**). While single Pp*cus1* and Pp*cus2* mutants were indistinguishable from wild-type plants, double Pp*cus1*/Pp*cus2* mutants displayed clear defects in gametophore development, characterized by reduced phyllid size and pronounced cellular disorganization (**Fig. 2C-E**).

**Figure 2.**
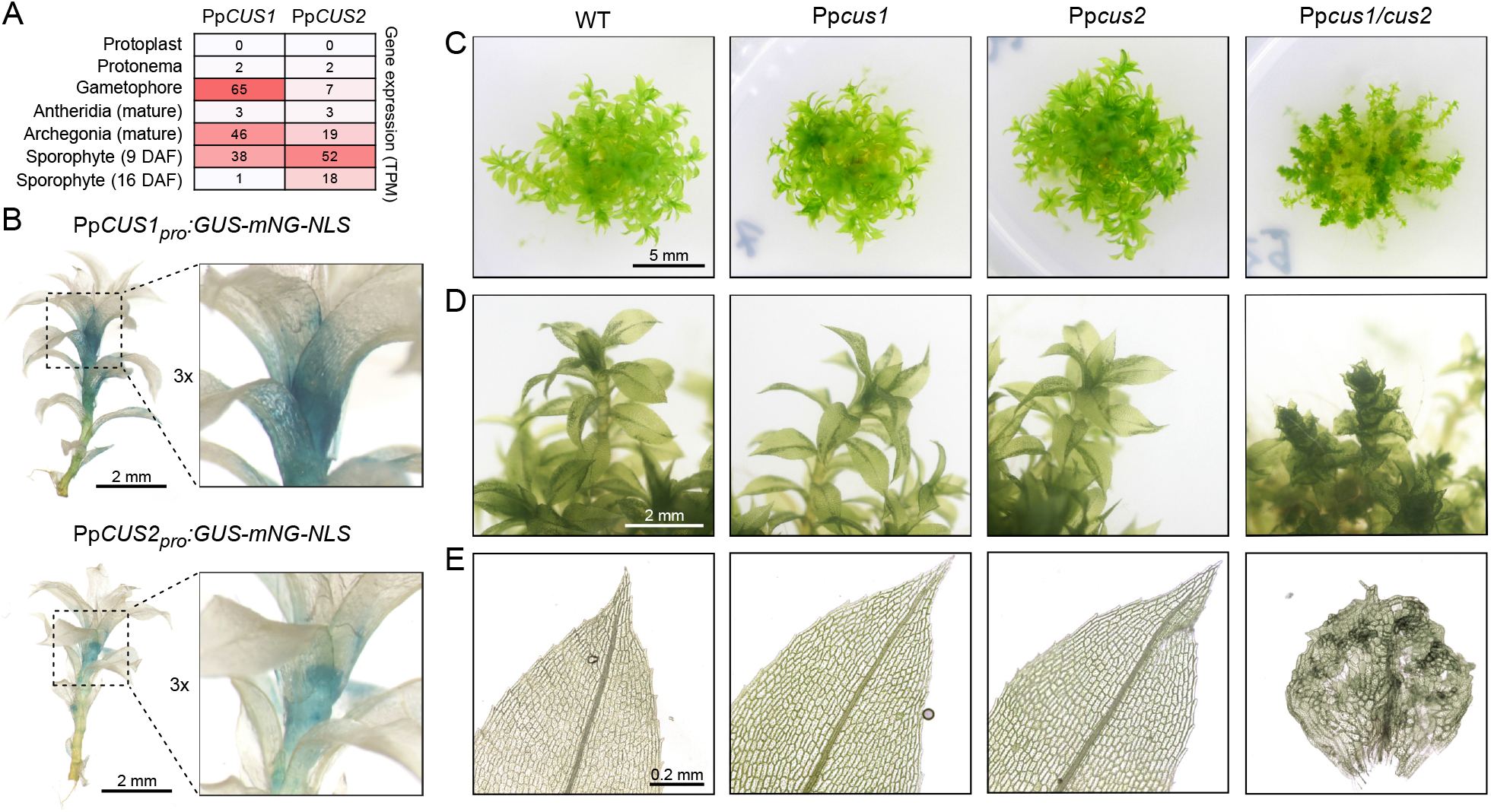
Inactivation of *P. patens CUS* genes impairs gametophore development. (**A**) Expression levels of the two *P. patens CU5* paralogs in various tissues, expressed as transcripts per kilobase million (TPM). Data are derived from the CoNekT database (Proost and Mutwil 2018). DAF, days after fertilization. (**B**) Representative patterns of Pp*CU51* and Pp*CU52* promoter activity in 8-week-old gametophores revealed by GUS staining (3h) of transcriptional reporter lines. A 3χ magnified view of the main site of expression of Pp*CU5* genes is shown. (**C-E**) Phenotypes of 8-week-old wild type, Pp*cus1* and Pp*cus2* single mutants, and the Pp*cus1/cus2* double mutant at the colony (**C**), gametophore (**D**), and phyllid (**E**) levels.

### *P. patens* CUS are apoplastic proteins essential for cuticle formation

Cutin polymerization is thought to occur locally in the apoplast according to angiosperm data (Girard et al. 2012; Yeats et al. 2012). We therefore examined whether the subcellular localization of *Pp*CUS proteins was consistent with this model. For both Pp*CUS* paralogs, we generated knock-in translational reporter lines in which the coding sequence of the mScarlet3 fluorophore (mSC3) was inserted between the region encoding the predicted signal peptide of *Pp*CUS proteins and the remainder of the coding sequence (**Supp. Fig. S4**). Each construct was individually integrated into the corresponding Pp*CUS* locus in a Pp*cus1/cus2* mutant background and successfully complemented the gametophore phenotype (**Fig. 3A; Supp. Fig. S4**), confirming that the chimeric mSC3-PpCUS proteins were functional. Confocal microscopy of phyllids revealed mSC3 fluorescence for both *Pp*CUS1 and *Pp*CUS2 at the cell periphery (**Fig. 3B**), consistent with an apoplastic localization. This was further confirmed by plasmolysis experiments, in which the mSC3 signal remained associated with the apoplast as the protoplast retracted (**Supp. Fig. S5**). We next examined cuticle structure in phyllids of Pp*cus1/cus2* double mutants by transmission electron microscopy. Electron micrographs revealed that cuticle integrity was compromised in the double mutant compared with wild-type plants (**Fig. 3C**), indicating a function for *Pp*CUS proteins in cuticle formation. Consistent with this defect, toluidine blue assays showed that Pp*cus1/cus2* gametophores were significantly more permeable (**Fig. 3D-E**), pointing to an alteration of the diffusion-limiting properties of the cuticle.

**Figure 3.**
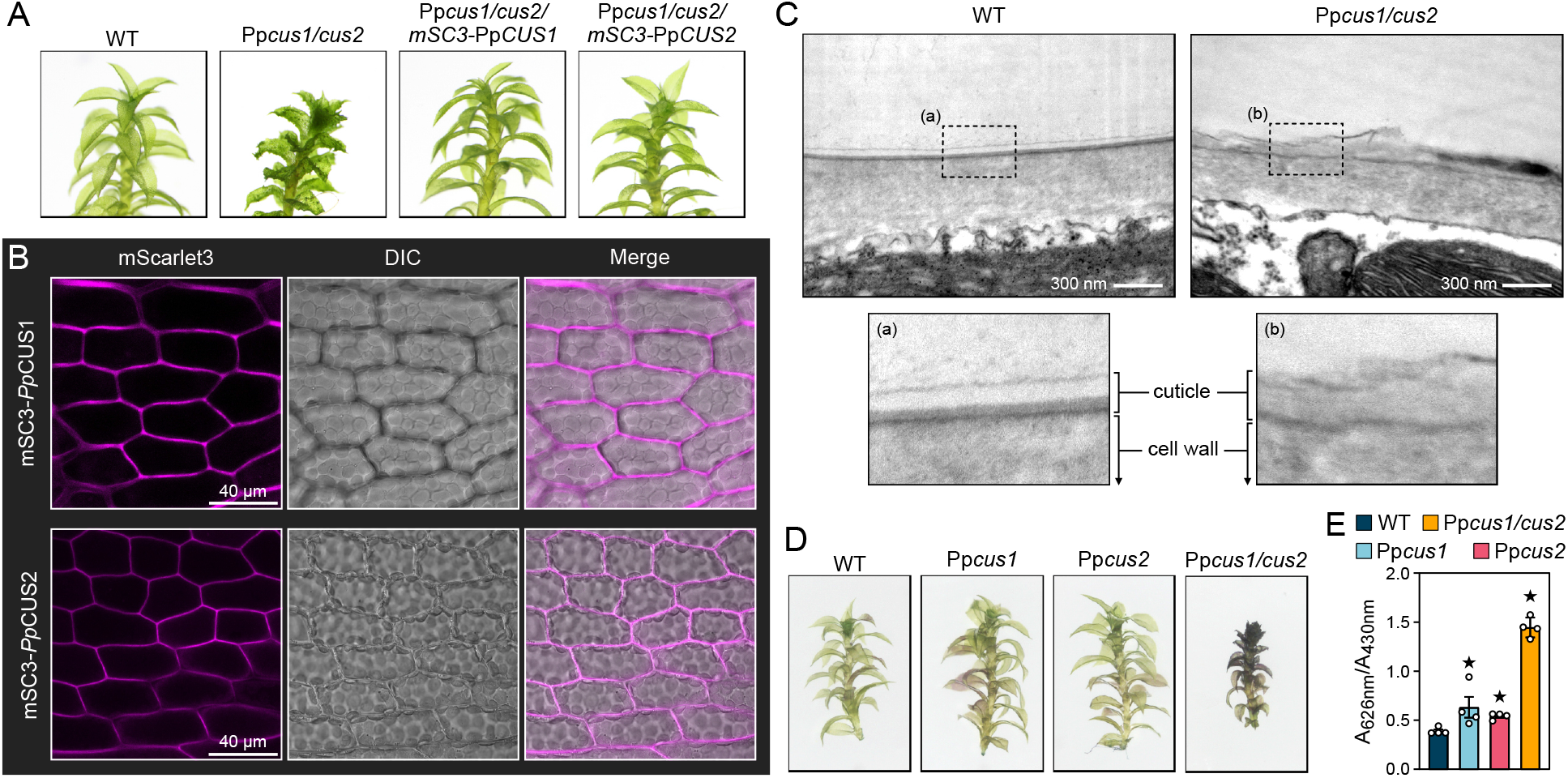
*Pp*CUS proteins localize to the apoplast and control cuticle integrity. (**A**) Phenotypes of 8-week-old gametophores from wild type, Pp*cus1/cus2*, and Pp*cus1/cus2* complemented with *m5C3-PpCU5* knock-in constructs. (**B**) Determination of mSC3-PpCUS protein localization by confocal microscopy in phyllids of 8-week-old Pp*cus1/cus2* complemented lines. (**C**) Transmission electron micrographs of wild-type and Pp*cus1/cus2* phyllid cross sections. Close-ups on the cuticle region are shown. (**D-E**) Assessment of cuticle permeability in 8-week-old gametophores with toluidine blue. (**D**) Pictures of gametophores after toluidine blue assay. (**E**) Quantification of toluidine blue in gametophores. Results are the means ± SEM of four biological replicates for WT and four independent lines for mutants. A star indicates a significant difference with WT according to Mann-Whitney *U* test (*p*<0.05).

### P. patens CUS control cuticle lipid polyester formation in planta

To further assess functional conservation within the CUS family, we examined the metabolic consequences of CUS deficiency in *P. patens*. We first analyzed 2-MHG, the substrate of tomato *Sl*CUS1 (Yeats et al. 2012) (**Fig. 4A**), in gametophore extracts of Pp*cus* mutants using LC-MS/MS. This analysis revealed a 10-fold accumulation of 2-MHG in Pp*cus1/cus2* double mutants compared with wild type (**Fig. 4A-B**), indicating that *Pp*CUS proteins employ the same substrate as their angiosperm homologs *in planta*. Accumulation of 2-MHG was also observed in Pp*cus1* single mutants, albeit to a lesser extent, consistent with Pp*CUS1* being the most expressed paralog in gametophores (**Fig. 2A**). We then examined the structural determinants of 2-MHG binding in CUS proteins using *Pp*CUS1 as a model. Docking analyses predicted that SER44, GLY120, ASN176, and ASN180 form hydrogen bonds with the glycerol moiety of 2-MHG (**Fig. 4C; Supp. Fig. S6A**). These interactions positioned the ester bond in close proximity to the catalytic triad SER44-ASP335-HIS338, consistent with the polymerization activity of CUS proteins involving 2-MHG hydrolysis (Yeats et al. 2012; Philippe et al. 2025). Moreover, 2-MHG ester bond was predicted to lie at the entrance of a surface-exposed protein tunnel (**Supp. Fig. S6B**), which would favor the incorporation of 10,16-diOH-C16:0 into the polymer.

**Figure 4.**
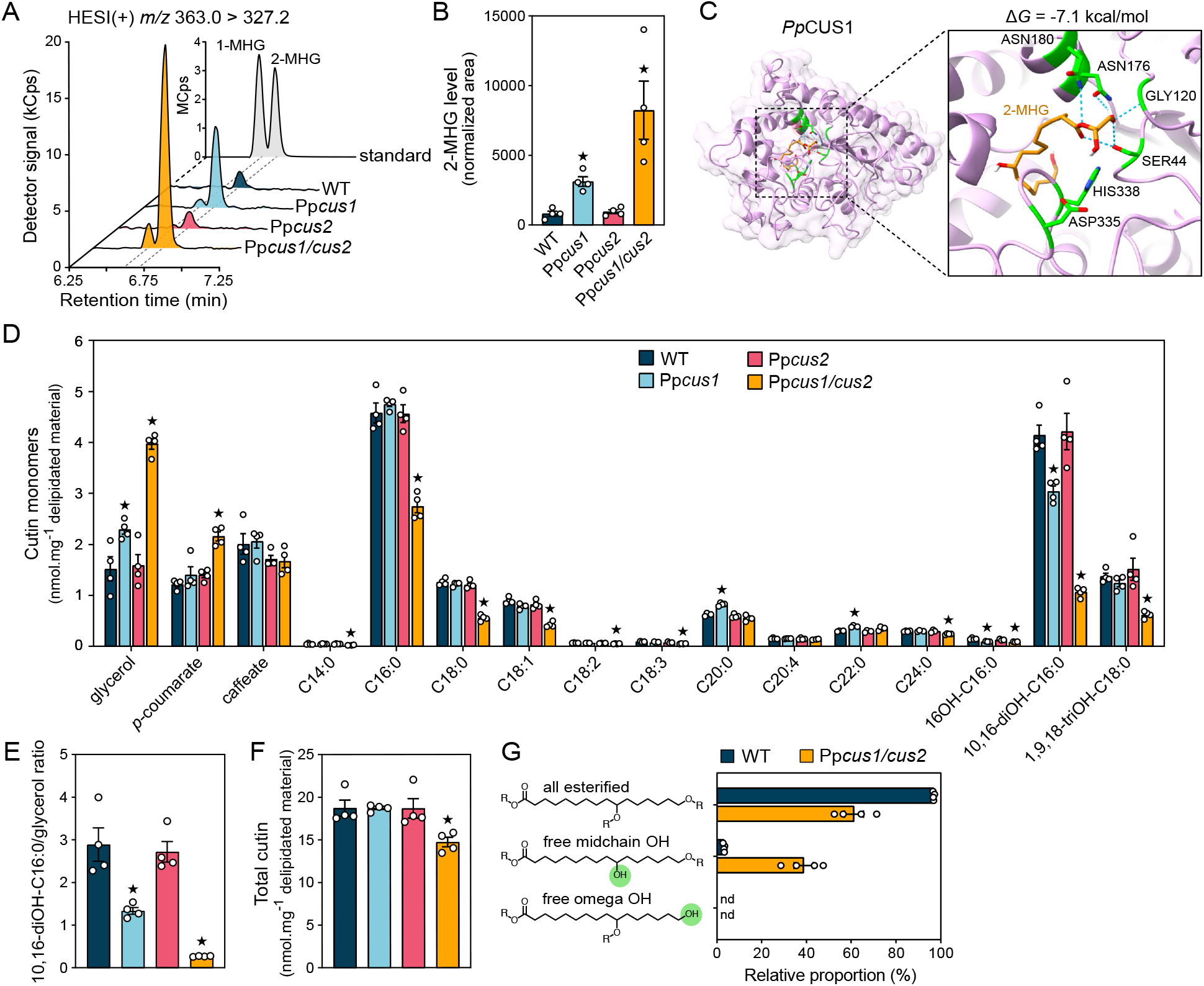
Pp*cus* mutants are defective in l0,l6-diOH-Cl6:0 polymerization. (**A-B**) Determination of 2-MHG levels in 8-week-old gametophores by UHPLC-MS/MS. (**A**) Chromatograms of metabolic extracts showing the 2-MHG diagnostic ion transition *m/z* 363.0 > 327.2. chromatogram of a 5 μM 2-MHG standard solution (courtesy of Prof. Jocelyn Rose, Cornell University) is also shown. Note that 2-MHG readily isomerizes into the thermodynamically favored 1-mono(10,16-dihydroxyhexadecanoyl)glycerol (1-MHG) (Yeats et al. 2012); consequently, both isomers are detected. (**B**) Relative quantification of 2-MHG levels. Peak area was normalized to plant dry weight. (**C**) Three-dimensional structure of the mature *Pp*CUS1 protein (residues 31-370) in complex with 2-MHG (orange). The docking pose corresponds to the state with the lowest Gibbs free energy (-7.1 kcal/mol). A close-up view highlights SER44, GLY120, ASN176, and ASN180 residues (green), which are predicted to form hydrogen bonds (blue dotted lines) with 2-MHG. In addition to SER44, the other two residues of the GDSL catalytic triad, ASP335 and HIS338, are also displayed (green). (**D-F**) Characterization of the cutin polymer from 8-week-old gametophores by chemical depolymerization followed with GC-TOFMS analysis. (**D**) Compositional analysis of the cutin polymer. (**E**) Total amount of cutin monomers. (**F**) Molar ratio of 10,16-diOH-C16:0 to glycerol at the whole cutin level. (**G**) Esterification levels of the different hydroxyl groups (OH) in 10,16-diOH-C16:0 cutin monomer from 8-week-old gametophores. Results in panels **B, D, E, F** and **G** are the means ± SEM of four biological replicates for WT and four independent lines for mutants. A star indicates a significant difference with WT according to Mann-Whitney *U* test (*p*<0.05).

Next, we analyzed cutin composition in wild-type and mutant gametophores using base-catalyzed methanolysis followed by GC-TOFMS determination of the released monomers. This analysis revealed a 75% reduction of 10,16-diOH-C16:0 in Pp*cus1/cus2* double mutants (**Fig. 4D**). Glycerol levels increased concomitantly in the double mutant, shifting the 10,16-diOH-C16:0-to-glycerol molar ratio from 2.9 in wild type to 0.27 in Pp*cus1/cus2* mutants at the whole cutin level (**Fig. 4E**). These observations support a role for *Pp*CUS proteins in producing a 10,16-diOH-C16:0 polyester from the 2-MHG precursor *in planta*, consistent with earlier in vitro data (Yeats et al. 2014). Overall, deficiency in both Pp*CUS* paralogs resulted in a 20% reduction in total cutin content (**Fig. 4F**). Finally, we investigated whether *Pp*CUS proteins, like angiosperm homologs (Philippe et al. 2016), are able to polymerize 10,16-diOH-C16:0 on both terminal and midchain hydroxyl groups (OH). To this end, free OH groups in cutin were benzyl-etherified prior to depolymerization and GC-MS analysis, providing a snapshot of the native esterification state of the 10,16-diOH-C16:0 monomer. In wild-type plants, 97% of the 10,16-diOH-C16:0 monomer were esterified at both the midchain and terminal OH groups, with only 3% having a free midchain OH (**Fig. 4G**). Inactivation of Pp*CUS* genes caused a sharp increase in 10,16-diOH-C16:0 monomers with a free midchain OH, reaching *ca*. 40% in Pp*cus1/cus2* mutants (**Fig. 4G**). Together, these results indicate that *Pp*CUS proteins can acylate both terminal and midchain OH groups, thus producing a reticulated 10,16-diOH-C16:0 polyester *in planta*.

## DISCUSSION

Although the cuticle is regarded as a hallmark of land plants, its evolutionary origins remain unclear. Our study shows that the CUS enzyme family, which plays a central role in cuticle formation in angiosperms (Girard et al. 2012; Yeats et al. 2012; Hong et al. 2017; Philippe et al. 2025), evolved during terrestrialization in a common ancestor of bryophytes and tracheophytes (**Fig. 1**), the two sister lineages of land plants (Morris et al. 2018; Puttick et al. 2018). Previous phylogenomic analyses reported that the *CYP, GPAT*, and *LACS* gene families involved in cutin precursor production evolved in parallel with, or possibly even before, the *CUS* family, suggesting that the complete cutin biosynthetic machinery was already established in the last common ancestor of embryophytes (Kong et al. 2020; Philippe et al. 2020). Yet, as the enzymes responsible for the final assembly of these precursors, CUS likely played a central role in the emergence of the cuticle by controlling the formation of its polymeric framework.

This view is supported by our data showing that CUS enzymes are essential to cuticle formation in the moss *P. patens*, catalyzing the synthesis of an apoplastic, reticulated 10,16-diOH-C16:0 polyester using 2-mono-(10,16-dihydroxyhexadecanoyl)glycerol as substrate (**Fig. 4**). This polymerization activity is consistent with that reported for CUS homologs in angiosperms (Girard et al. 2012; Yeats et al. 2012; Philippe et al. 2025), pointing to a remarkable conservation of CUS catalytic characteristics over 500 million years of embryophyte evolution. Inactivation of *CUS* genes compromised moss cuticle integrity, causing a dramatic increase in tissue permeability (**Fig. 3**), which is in line with the phenotypes observed in tomato fruits and Arabidopsis flowers (Girard et al. 2012; Yeats et al. 2012; Hong et al. 2017; Philippe et al. 2025). Together, these findings delineate a cornerstone and deeply conserved role for CUS enzymes in shaping the cuticle and controlling its diffusion-limiting properties in land plants. In this context, it is reasonable to propose that the emergence of the CUS enzyme family in an ancestral embryophyte represented a key biochemical innovation that facilitated plant colonization of terrestrial habitats.

Our study also highlights a developmental role for CUS, and thus for the cuticle, that extends beyond protective functions. Phyllids of CUS-deficient moss mutants displayed altered cell shape (**Fig. 2**). This phenotype echoes with recent studies showing that the cuticle contributes to the control of anisotropic cell growth in both *P. patens* and Arabidopsis (Zhang et al. 2024; Raggi et al. 2026). Whether this function underlies the abnormal phyllid development observed in moss remains an open question. These observations complement earlier work reporting that a functional cuticle is required during organogenesis to prevent organ fusion in *P. patens* and Arabidopsis (Wellesen et al. 2001; Renault et al. 2017). Collectively, our results together with previous studies indicate that the cuticle played an ancestral developmental role in addition to its protective functions, a dual role that likely contributed to its strict conservation across land plants.

## METHODS

### Phylogenetic reconstruction

Homologs of the functionally characterized *Solanum lycopersicum* CUS1 protein (*Solyc11g006250*) (Girard et al. 2012; Yeats et al. 2012) were searched with phmmer v3.4 (http://hmmer.org) in 23 Viridiplantae genomes (**Supp. Fig. S1**). Sequence retrieval was limited to homologs with an E-value < 1e^-20^ and a minimum length of 320 amino acids, yielding a dataset of 610 proteins. Sequences were aligned using MUSCLE v3.8.31 (Edgar 2004) prior to maximum-likelihood phylogenetic reconstruction with IQ-TREE v3.0.1 (Wong et al. 2025). The best-fitting protein evolutionary model was automatically selected using ModelFinder (Kalyaanamoorthy et al. 2017) based on Bayesian Information Criterion. Branch support was assessed with 10,000 ultrafast bootstrap replicates (Minh et al. 2013). The resulting tree was visualized using the ggtree R package (Yu et al. 2017) and further annotated with Affinity Designer.

### Plant material and growth conditions

The Reute strain of *Physcomitrium patens* (Hiss et al. 2017) was obtained from the International Moss Stock Center (catalog n°40040) and used as the wild type for all experiments. *P. patens* was grown axenically on solid Knop medium under a 16h light photoperiod at 22°C/18°C (day/night), with a light intensity of 50 µmol.m^-2^.s^-1^ and 60% relative humidity. Knop medium contained 250 mg/L MgSO_4_ · 7H_2_O, 250 mg/L KH_2_PO_4_, 250 mg/L KCl, 1000 mg/L Ca(NO_3_)_2_ · 4H_2_O, 12.5 mg/L, FeSO_4_ · 7H_2_O, 0.012 mg/L CoCl_2_ · 6H_2_O, 0.012 mg/L CuSO_4_ · 5H_2_O, 3.09 mg/L H_3_BO_3_, 0.42 mg/L KI, 8.5 mg/L MnSO_4_ · H_2_O, 0.12 mg/L Na_2_MoO_4_ · 2H_2_O and 4.3 mg/L ZnSO_4_ · 7H_2_O. The medium was adjusted to pH 5.8 with KOH and solidified with 1.2% agar for plate cultures. Protonema was cultured in 500 mL flasks filled with 200 mL liquid Knop medium constantly agitated at 130 rpm, and maintained by weekly tissue homogenization using an IKA ULTRA-TURRAX T25 equipped with S25N-18G dispersing tool. Gametophore liquid cultures were initiated by soft disruption of gametophores grown on Knop solid medium.

### CRISPR/Cas9-mediated gene inactivation

*P. patens* gene knock-outs were generated by transient CRISPR/Cas9 expression according to Lopez-Obando et al. (Lopez-Obando et al. 2016), with some modifications. A 20 nucleotide protospacer sequence located in the first exon of each Pp*CUS* gene (**Supp. Fig. S3**) was identified using the online CRISPOR tool (Concordet and Haeussler 2018). Complementary oligonucleotides were annealed to reconstitute double-stranded protospacers (**primers in Supp. Tab. S1**), which were cloned into the *pENTR-PpU6-sgRNA-L1L2* vector (Mallett et al. 2019) using *Bsa*I restriction enzyme, enabling sgRNA expression under control of Pp*U6* promoter. *P. patens* protoplasts were transfected with 5 µg of *pENTR-PpU6-sgRNA*, 10 µg *pACT-Cas9* (SpCas9 expression) (Lopez-Obando et al. 2016), and 5 µg *pRT101-NPTII* (transformant selection) (Girke et al. 1998) via a polyethylene glycol-mediated protocol (Hohe et al. 2004). For the double mutants, protoplasts were simultaneously transfected with 5 µg of each *pENTR-PpU6-sgRNA* plasmid. After 10 days of regeneration, plants underwent a single round of selection on solid Knop medium supplemented with 25 µg/mL G418 for two weeks. Resistant plants were transferred to standard Knop plates and allowed to grow further before genotyping. Mutations in individual moss lines were verified by Sanger sequencing of ∼600 bp amplicons generated using Phusion High-Fidelity DNA polymerase and primers spanning target sites (**Supp. Fig. S3, primers in Supp. Tab. S1**). For each genotype, four independent lines carrying a frameshift mutation were selected and used as experimental replicates (**Supp. Fig. S3**).

### CRISPR/Cas9-assisted gene knock-in

Targeted knock-in into the *P. patens* genome was achieved by homologous recombination (HR) enhanced by transient CRISPR/Cas9 expression. To this end, the DNA construct of interest was assembled by Gibson cloning together with two flanking DNA fragments (750-1001 bp) homologous to the target genomic region into a pGEM-T-Easy backbone, generating HR donor vectors. For Pp*CUSpro:GUS-mNG-NLS* reporter lines, constructs were inserted into the *PTA2* neutral locus (Kubo et al. 2013) of wild-type plants using a sgRNA targeting *PTA2* 3’ UTR (**Supp. Fig. S2**). For *mSC3-*Pp*CUS* complemented lines, constructs were inserted into the Pp*CUS* loci of the *Ppcus1/cus2* #52 line using sgRNAs targeting the first intron of Pp*CUS* genes (**Supp. Fig. S4**). Transformants were produced by transfecting *P. patens* protoplasts with 5 µg of *pENTR-PpU6-sgRNA*, 10 µg *pACT-Cas9* (Lopez-Obando et al. 2016), 5 µg *pRT101-NPTII* (Girke et al. 1998), and 10 µg of HR donor plasmid *via* a polyethylene glycol-mediated protocol (Hohe et al. 2004). Selection was performed with G418 as described above. Correct integration of the DNA constructs at both the 5′ and 3′ ends was verified by PCR using genomic DNA from individual transformants and specific primer pairs (**Supp. Fig. S2, S4 and Supp. Tab. S1**).

### GUS staining

Plant tissues were vacuum infiltrated for 10 min with X-Gluc solution (50 mM potassium phosphate buffer pH 7.0, 0.5 mM ferrocyanide, 0.5 mM ferricyanide, 0.1% Triton X-100, 0.5 mg/mL X-Gluc) and incubated at 37°C in the dark. Chlorophyll was removed by washing tissues three times in 70% ethanol. Pictures were taken with an Olympus SZX7 stereomicroscope mounted with a Canon EOS R8 Camera *via* a LMscope DD2XZ42 adapter.

### Toluidine blue assay

*P. patens* tissue permeability was assessed by immersing gametophores in toluidine blue solution as previously described (Knosp et al. 2024).

### Laser-scanning confocal microcopy

Localization of mSC3-*Pp*CUS proteins in 8-week-old moss gametophores was performed using an inverted Zeiss LSM 980 confocal microscope with a 63× Plan-Apochromat objective (numerical aperture 1.4, oil immersion). mScarlet3 was excited at 543 nm, and emission was collected from 582-602 nm. To confirm apoplastic localization, gametophores were incubated in 0.4 M mannitol for 30 minutes to induce plasmolysis prior to imaging. Image processing was carried out using Fiji (Schindelin et al. 2012).

### Transmission electron microscopy

Samples were fixed overnight in 2% glutaraldehyde, post-fixed with 0.5% (v/v) osmium tetroxide in 150 mM phosphate buffer (pH 7.2) for 2h, and stained overnight with 2% (w/v) uranyl acetate. After dehydration through a graded ethanol series, samples were infiltrated with EPON812 resin (Polysciences) and polymerized for 48h at 60°C. Ultrathin sections (70 nm) were cut with an Ultracut E microtome (Reichert), collected on formvar-coated grids (Electron Microscopy Sciences), and imaged with a Zeiss Sigma 300HV electron microscope at 30 kV.

### 2-MHG profiling in plant extracts

Eight-week-old gametophores grown in liquid culture were collected, briefly blotted on paper towel, and snap-frozen in liquid nitrogen. Samples were lyophilized overnight and homogenized using 5 mm steel beads in a Tissuelyser II (Qiagen) for 1 min at 30 Hz. 2-MHG was extracted from 8 mg of dry plant powder using a methanol:chloroform:water protocol as described in (Knosp et al. 2024). Crude extracts were dried under vacuum and concentrated tenfold in 50% methanol prior to analysis. Metabolite separation and detection were performed using a Dionex UltiMate 3000 (ThermoFisher Scientific) UHPLC coupled to an EvoQ Elite LC-TQ (Bruker) triple quadrupole mass spectrometer, as previously reported (Kriegshauser et al. 2021). A 5 µL aliquot was injected onto a C18 Cortecs UPLC HSS T3 column (150 × 2.1 mm, 1.6 µm; Waters) maintained at 35°C, and eluted with LC-MS-grade water (A) and acetonitrile (B), both containing 0.1% formic acid, at a flow rate of 0.35 mL.min^-1^ using the following gradient: 0.0 min, 5% B; 6.0 min, 50% B; 16.0 min, 100% B; 18.0 min, 100% B; 18.5 min, 5% B; 20.0 min, 5% B. 2-MHG was ionized in positive mode using a heated electrospray ionization source (HESI, Bruker) and detected via the specific *m/z* 363.0 > 345.2 transition (collision energy of 6 eV), established using an authentic 2-MHG standard (courtesy of Prof. Jocelyn K.C. Rose, Cornell University). Bruker MS Data Review software was used to integrate peaks and report corresponding areas. 2-MHG levels were expressed in relative levels, with peak areas normalized to the dry weight of the extracted plant material.

### Compositional analysis of cutin

Chemical depolymerization of *P. patens* cutin was performed using a base-catalyzed protocol as described in (Philippe et al. 2016), except that heptadecanoic acid (C17:0) and ribitol were used as internal standards. Released cutin monomers, including glycerol, were analyzed as described in (Knosp et al. 2024) using gas chromatography coupled to a time-of-flight mass spectrometer (Pegasus BT2, LECO) and quantified based on analyte-specific response factors (**Supp. Tab. S3**).

### Chemical labeling of cutin free hydroxyl groups

Free hydroxyl groups within the cutin polymer were derivatized *via* benzyl etherification and analyzed by GC-MS following a protocol described in (Philippe et al. 2016).

### Docking experiments

The AlphaFold-predicted three-dimensional (3D) structure of *Pp*CUS1 was retrieved from the UniProt database (accession no. A9RXD4). Prior to docking, the molecular geometry of the 2-MHG ligand was optimized using Avogadro with the Universal Force Field approach and the steepest descent algorithm (Hanwell et al. 2012). 2-MHG was then docked into the *Pp*CUS1 3D model lacking the signal peptide (residues 31-370) using AutoDock Vina v1.1.2 in rigid mode (Trott and Olson 2010) with a 40 Å × 40 Å × 40 Å search box. Ligand and receptor files required for docking experiments were prepared with AutodockTools 1.5.7 (https://ccsb.scripps.edu/mgltools/). Docking results were visualized using ChimeraX (Pettersen et al. 2021).

## ACCESSION NUMBERS

Pp*CUS1, Pp3c14_19530V3*.*1*; Pp*CUS2, Pp3c17_21660V3*.*1*; Mp*CUS1, Mp8g13940*.*1*; Aa*CUS1, AagrBONN_117*.*3162*.*2*; Sk*CUS1, SK07G10970*.*1*; Cr*CUS1, Ceric*.*10G057300*.*1*; Gb*CUS1, Gbiloba*.*chr9*.*769*; Sl*CUS1, Solyc11g006250*.*2*.*1*; At*CUS2, AT5G33370*.*1*

## AUTHOR CONTRIBUTIONS

Conceptualization: SK, HR

Methodology: SK, BB

Investigation: SK, FB, EL, GZ, LM, ME, BB, HR

Visualization and Writing—original draft: HR

Writing—review & editing: SK, FB, EL, GZ, LM, ME, BB, HR

Supervision: HR

## COMPETING INTEREST STATEMENT

Authors declare no competing interests.

## ACKNOWLEDGMENTS

The authors thank Prof. Jocelyn K.C. Rose (Cornell University) for providing the authentic 2-MHG standard, and Dr. Fabien Nogué (INRAE, France) for providing pAct-Cas9 vector. HR received support from the Initiative of Excellence IDEX University of Strasbourg and the Agence Nationale de la Recherche (ANR-19-CE20-0017; ANR-24-CE20-0934). SK, FB, and EL were supported by PhD fellowships from the Ministère de l’Enseignement Supérieur, de la Recherche et de l’Espace. We also acknowledge the IBMP Metabolomics Analysis of Small-molecule Signatures (MASS) and Gene Expression Analysis core facilities for technical assistance.

## Notes

### Competing Interest Statement

The authors have declared no competing interest.

