## Supporting Materials for "The CUTIN SYNTHASE enzyme family was a key driver of cuticle emergence in land plants"

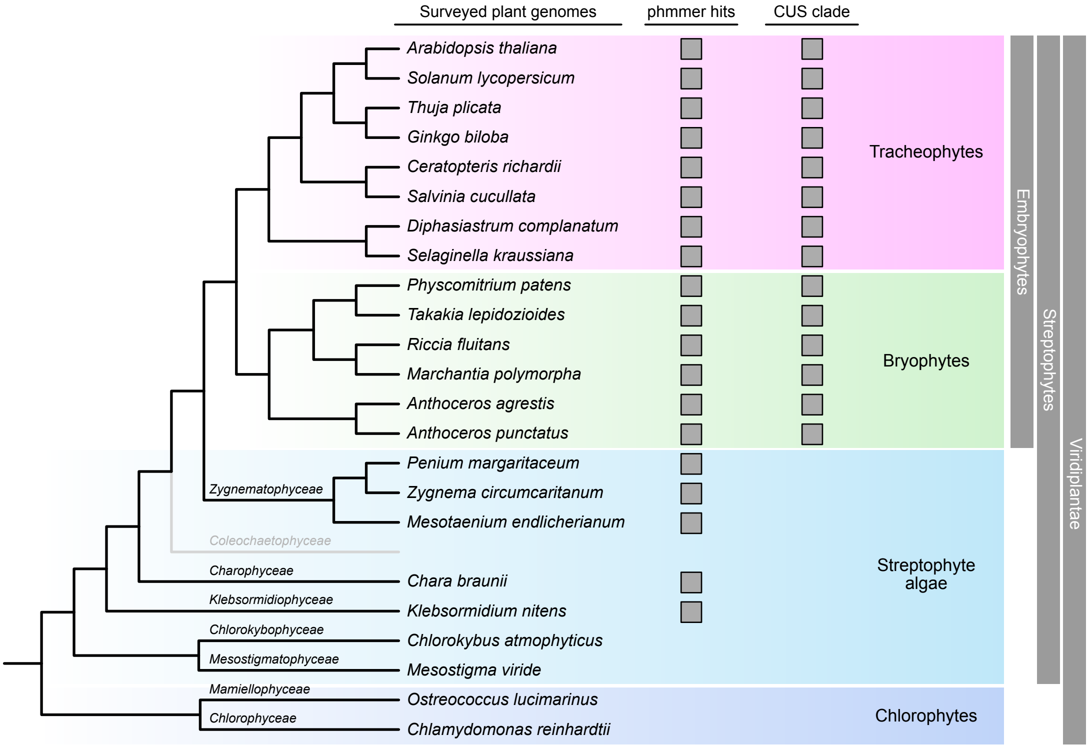


**Figure S1. Taxonomic sampling and overall output of the CUS homolog survey.**
Homologs of the tomato *Sl*CUS1 protein were searched using phmmer (E-value < e-20) in the following 23 Viridiplantae genomes: *Chlamydomonas reinhardtii* v5.5 (Merchant et al. 2007), *Ostreococcus lucimarinus* v2.0 (Palenik et al. 2007), *Chlorokybus atmophyticus* v1.0 (Wang et al. 2019), *Mesostigma viride* v1.0 (Wang et al. 2019), *Klebsormidium nitens* v1.1 (Hori et al. 2014), *Chara braunii* v1.0 (Nishiyama et al. 2018), *Mesotaenium endlicherianum* v2.0 (Dadras et al. 2023), *Zygnema circumcaritanum* v1.0 (Feng et al. 2024), *Penium margaritaceum* v1.0 (Jiao et al. 2020), *Anthoceros agrestis* (Bonn strain) v1.0 (Li et al. 2020), *Anthoceros punctatus* v1.0 (Li et al. 2020), *Physcomitrium patens* (Gransden strain) v3.3 (Lang et al. 2018), *Takakia lepidozioides* v1.0 (Hu et al. 2023), *Marchantia polymorpha* (Tak-1 strain) v7.1 (Tanizawa et al. 2025), *Riccia fluitans* v1.0 (bioproject PRJNA1158334), *Selaginella kraussiana* v2.0 (Liu et al. 2023), *Diphasiastrum complanatum* v3.1 (bioproject PRJNA914350), *Ceratopteris richardii* v2.1 (Marchant et al. 2022), *Salvinia cucullata* v1.2 (Li et al. 2018), *Ginkgo biloba* v2.0 (Liu et al. 2021), *Thuja plicata* v3.1 (Shalev et al. 2022), *Solanum lycopersicum* v4.0 (Hosmani et al. 2019) and *Arabidopsis thaliana* TAIR10 (Lamesch et al. 2012). Grey boxes indicate the presence of a homolog in the defined species.

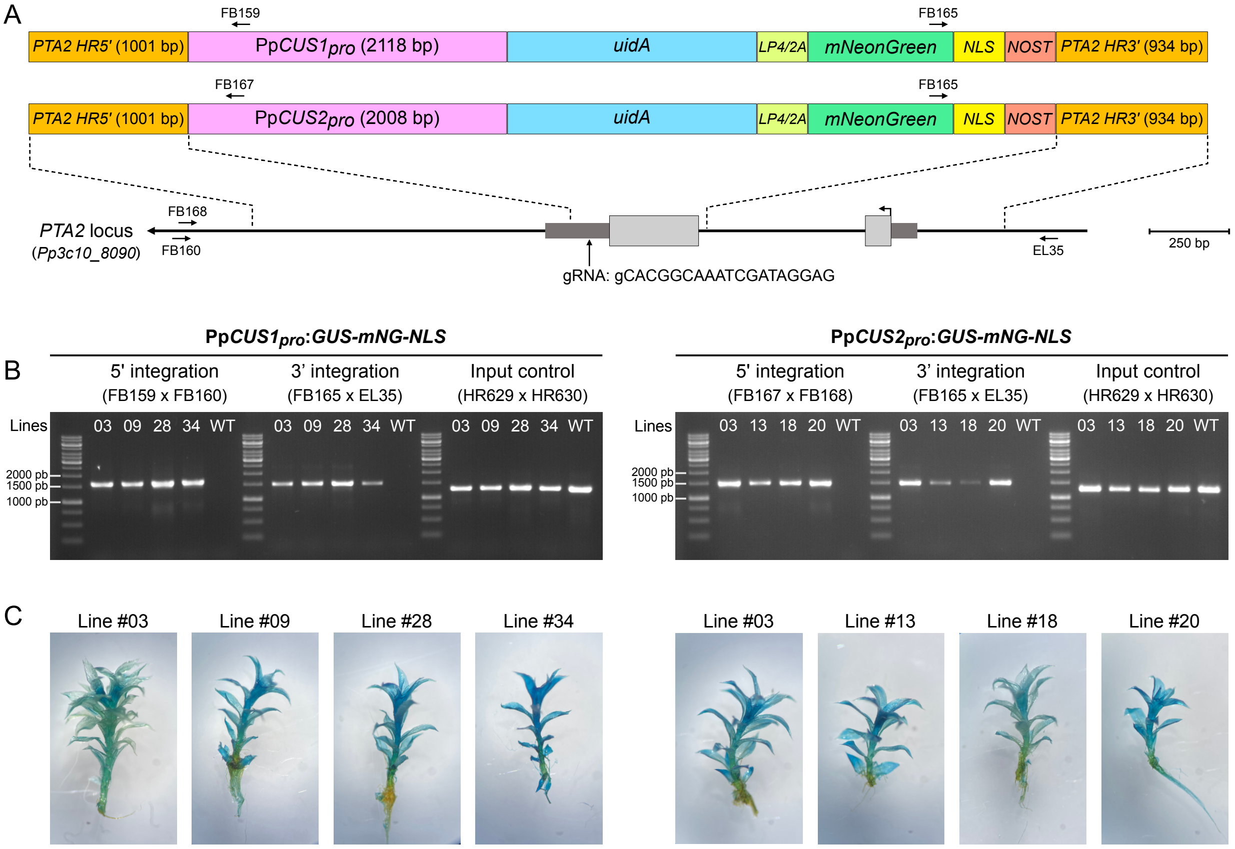


**Figure S2. Generation and validation of Pp*CUSpro:GUS-mNG-NLS* reporter lines.**
(**A**) Strategy for generating *Pp*CUS promoter-reporter lines. The promoter regions of Pp*CUS1* (2118 bp) and Pp*CUS2* (2008 bp) were PCR-amplified and assembled with a *uidA-mNeonGreen-NLS* reporter construct into a pGEM-T Easy vector containing *PTA2* homologous recombination fragments. The *uidA* and *mNeonGreen-NLS* coding sequences are fused *via* a hybrid *LP4/2A* linker sequence (François et al. 2004). The target site of the gRNA used for CRISPR-assisted homologous recombination is indicated. For proper expression under the Pp*U6* promoter, the first nucleotide of the protospacer was replaced by a guanine (in lowercase). Only the *PTA2* locus is drawn to scale. (**B**) PCR validation of correct construct integration at the *PTA2* genomic locus in G418-selected transgenic lines. Primer binding sites are indicated in (A). An unrelated genomic region was amplified as an input control. (**C**) GUS staining patterns after 8h of incubation in validated Pp*CUSpro:GUS-mNG-NLS* reporter lines.

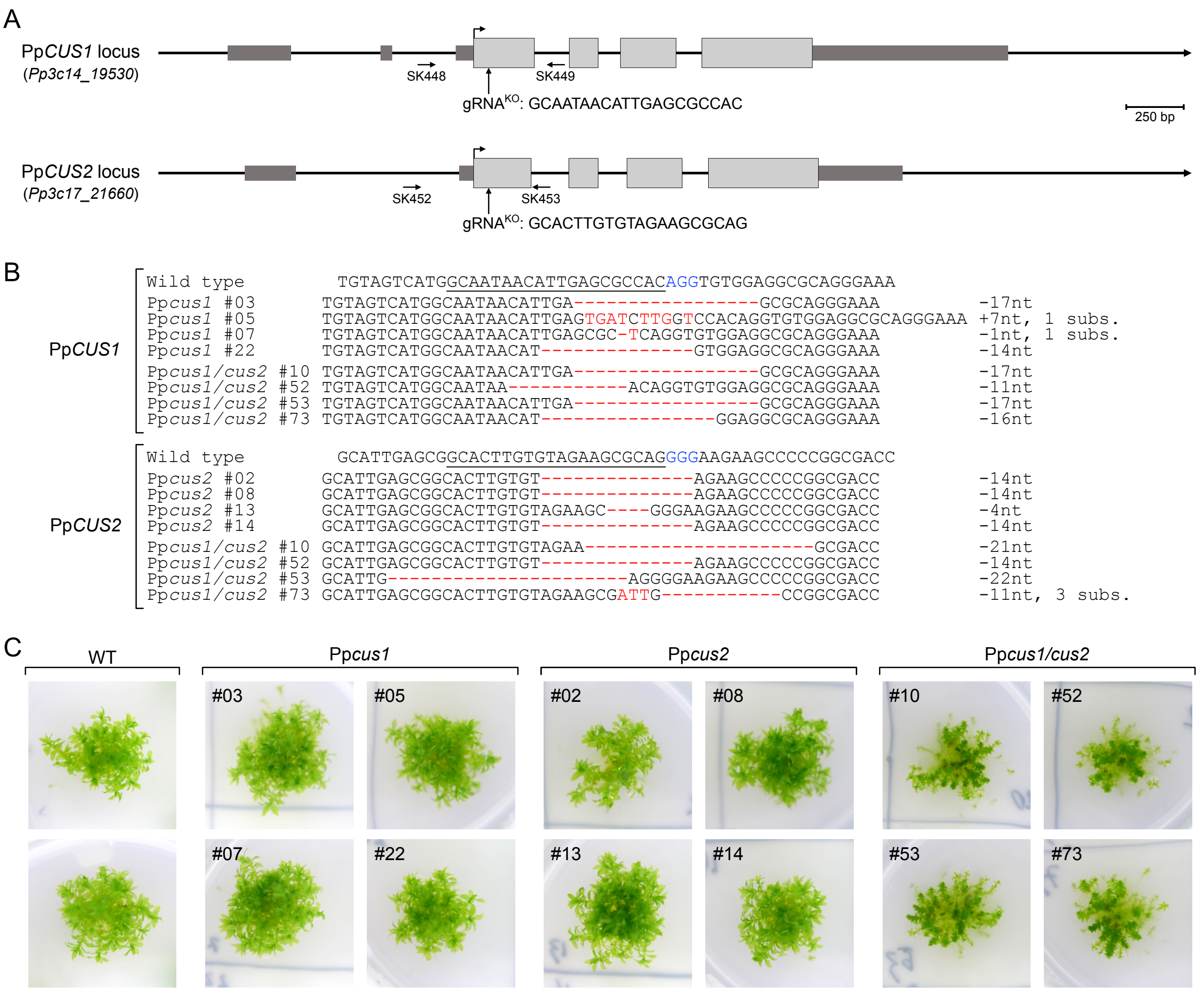


**Figure S3. Generation and validation of Pp*cus* null mutant lines.**
(**A**) Strategy for CRISPR/Cas9-mediated Pp*CUS* gene inactivation. CRISPR protospacer sequences and target sites are indicated. (**B**) Mutant alleles of G418-selected lines were characterized by Sanger sequencing of locus-specific PCR products; primer binding sites are indicated in (A). Nucleotide changes are highlighted in red. (**C**) Phenotypes of 8-week-old colonies from all validated lines used in the study.

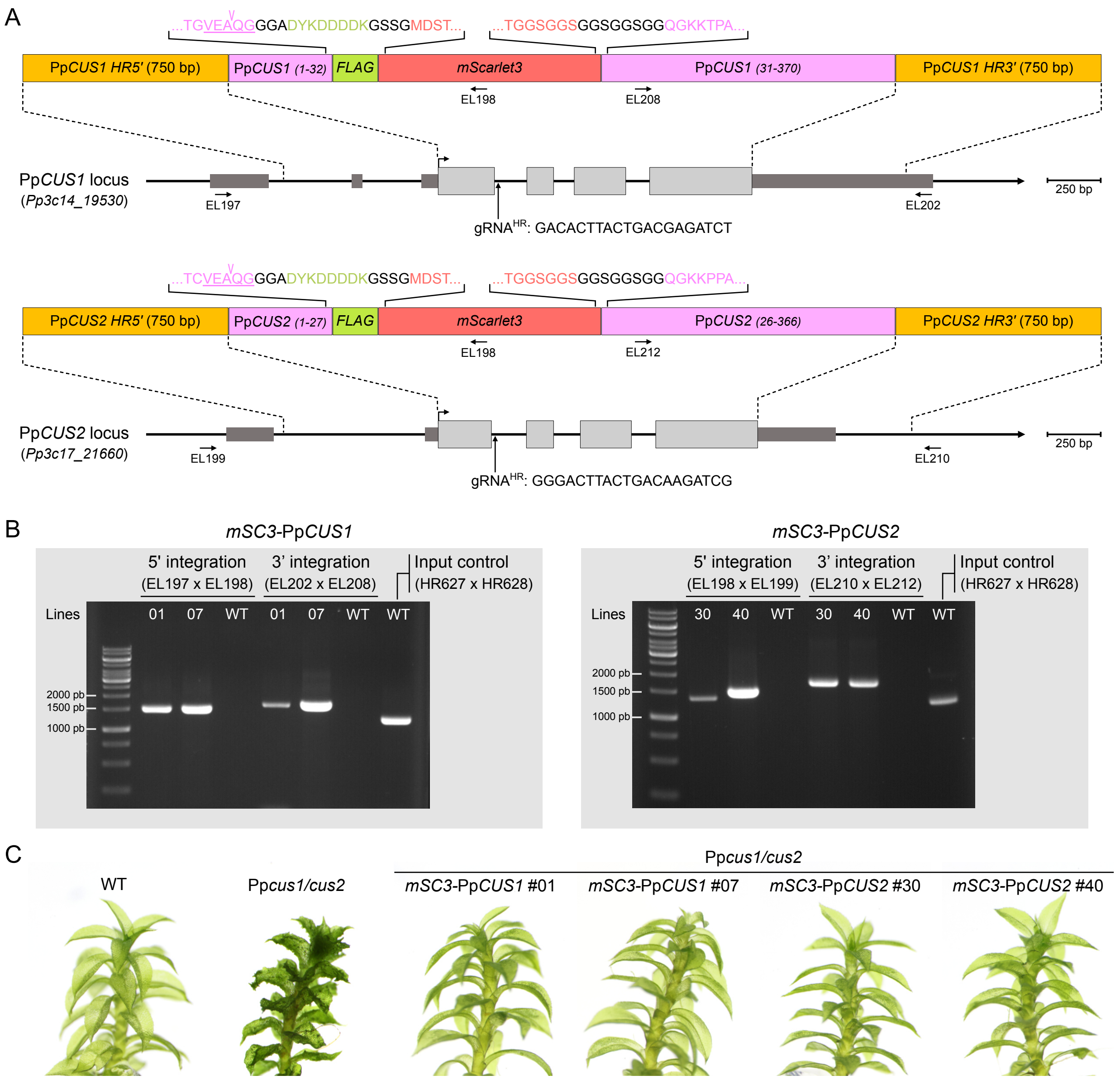


**Figure S4. Generation and validation of *mSC3*-Pp*CUS* complemented lines.**
(**A**) Strategy for the functional complementation of the Pp*cus1/cus2* mutant using CRISPR/Cas9-assisted gene knock-in. The *mScarlet3* coding sequence was inserted into *Pp*CUS coding sequences between the signal peptide (including the cleavage site; underlined) and the remainder of the protein (including the two amino acids immediately downstream of the cleavage site), resulting into the *mSC3*-Pp*CUS* chimeric construct. A FLAG tag (shown in green) was added to the *N*-terminus of mScarlet3. The amino acid sequences corresponding to the two transition regions between *Pp*CUS proteins and mScarlet3 are indicated. The genomic Pp*CUS* regions, from START to STOP codon, were replaced by *mSC3*-Pp*CUS* sequences via homologous recombination (HR) using 750 bp-flanking regions. HR efficiency was enhanced by inducing double-strand breaks within the first intron of *Pp*CUS genes using CRISPR/Cas. Corresponding protospacer sequences are shown. (**B**) PCR validation of correct construct integration of *mSC3*-Pp*CUS* constructs at the corresponding *Pp*CUS genomic locus in G418-selected transgenic lines. Primer binding sites are indicated in (A). An unrelated genomic region was amplified as an input control. (**C**) Macroscopic phenotypes of validated *mSC3*-Pp*CUS* complemented lines.

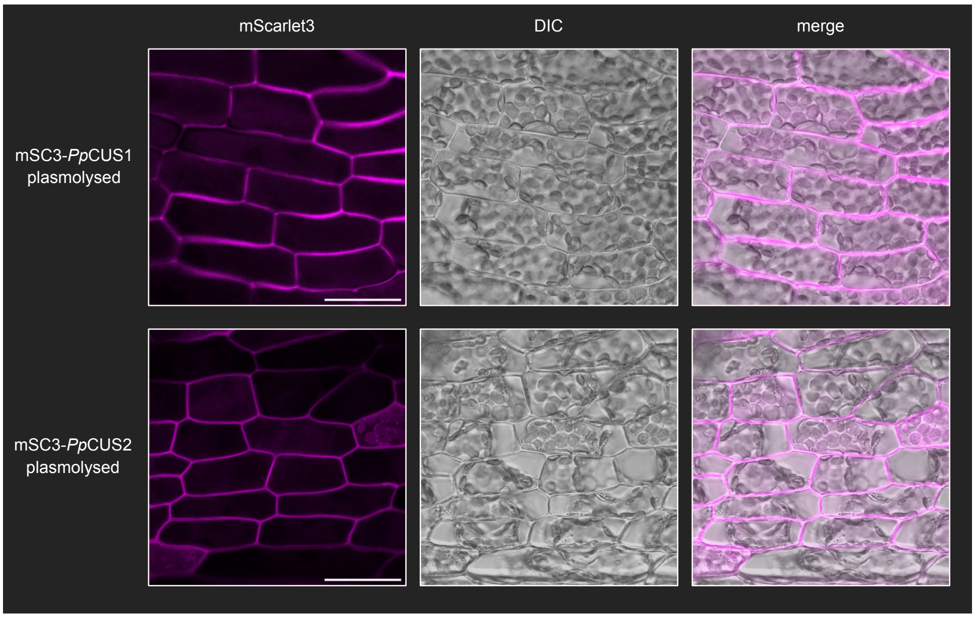


**Figure S5. *Pp*CUS proteins localize to the apoplast after plasmolysis.**
Localization of mSC3-*Pp*CUS protein was analyzed by confocal microscopy in phyllids of 8-week-old Pp*cus1/cus2* complemented lines incubated in 0.4 M mannitol for 30 min prior to imaging. Scale bars, 40 µm.

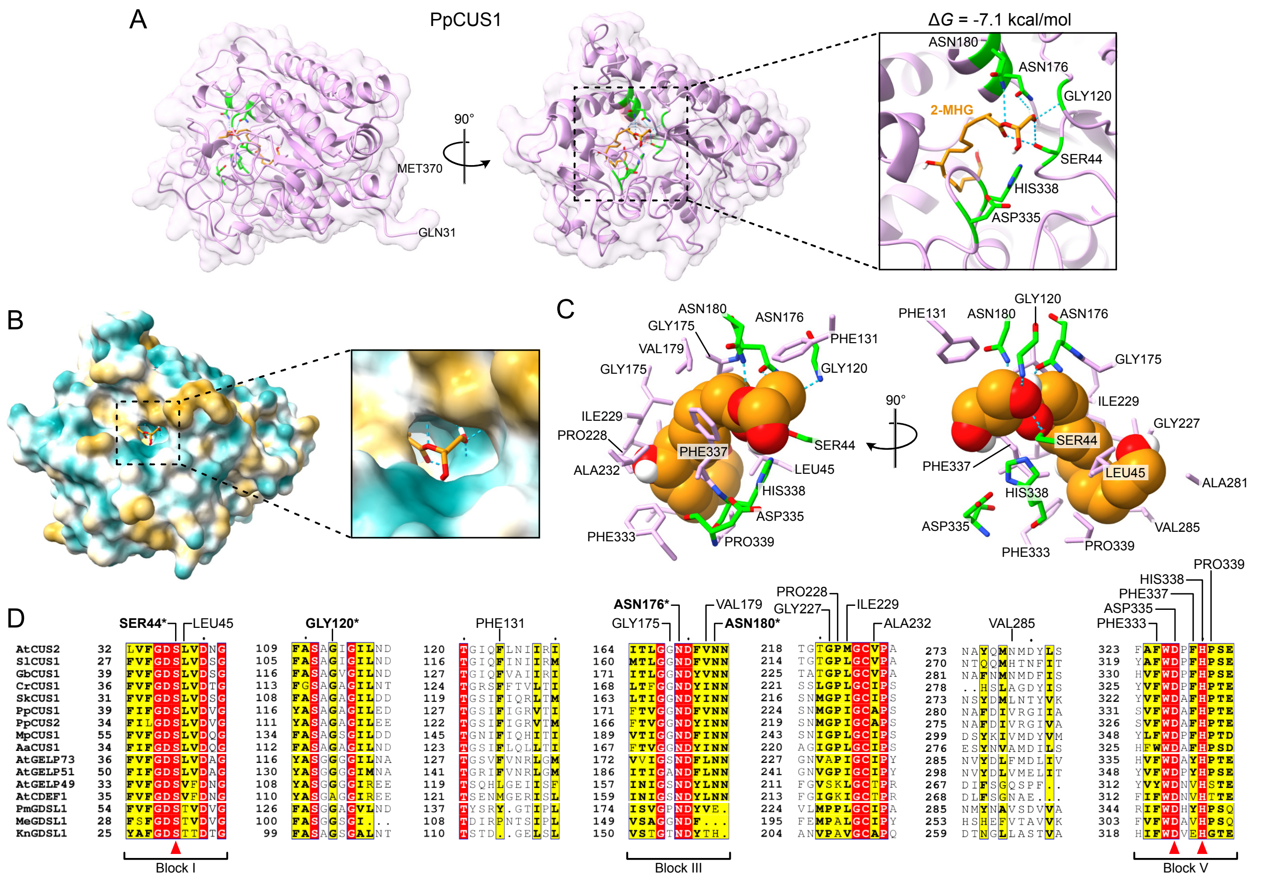


**Figure S6. Structural analysis of 2-MHG binding into CUS proteins.**
(**A**) Three-dimensional (3D) structure of the mature PpCUS1 protein (residues 31-370) in complex with 2-MHG (orange). The displayed pose corresponds to the state with the lowest Gibbs free energy (-7.1 kcal/mol). A close-up view highlights SER44, GLY120, ASN176, and ASN180 residues (green), which are predicted to form hydrogen bonds (blue dotted lines) with 2-MHG. In addition to SER44, the other two residues of the GDSL catalytic triad, ASP335 and HIS338, are also displayed (green). (**B**) Molecular hydrophobicity potential view of mature PpCUS1 protein in complex with 2-MHG. (**C**) 3D representation of *Pp*CUS1 residues making hydrophobic contacts with 2-MHG (orange). The residues forming hydrogen bonds and members of the catalytic triad are also shown in green. (**D**) Multiple sequence alignment of motifs containing PpCUS1 residues predicted to interact with 2-MHG. Representative CUS members from major embryophyte groups were aligned together with members of the CUS sister clade (*At*CDEF1, *At*GELP49, *At*GELP51, and *At*GELP73; see **Fig. 1B**). The best *Sl*CUS1 homologs from each of the streptophyte algae *Penium margaritaceum*, *Mesotaenium endlicherianum*, and *Klebsormidium nitens*, based on phmmer E-values, were also included in the alignment. Interacting residues were found in some of the conserved GDSL blocks defined in (Akoh et al. 2004). Red arrowheads indicate residues of the GDSL catalytic triad.

**Table S1. List of primers used in the study.**

| ID | Name | Sequence (5'>3') |
|---|---|---|
| **CRISPR protospacers (Golgen Gate cloning)** | | |
| SK0369 | PpPTA2_gRNA_KI_BsaI_F | ccatgCACGGCAAATCGATAGGAG |
| SK0370 | PpPTA2_gRNA_KI_BsaI_R | aaacCTCCTATCGATTTGCCGTGc |
| SK0446 | PpCUS1_gRNA_KO_BsaI_F | ccatGCAATAACATTGAGCGCCAC |
| SK0447 | PpCUS1_gRNA_KO_BsaI_R | aaacGTGGCGCTCAATGTTATTGC |
| SK0450 | PpCUS2_gRNA_KO_BsaI_F | ccatGCACTTGTGTAGAAGCGCAG |
| SK0451 | PpCUS2_gRNA_KO_BsaI_R | aaacCTGCGCTTCTACACAAGTGC |
| SK0609 | PpCUS1_gRNA_KI_BsaI_F | ccatGACACTTACTGACGAGATCT |
| SK0610 | PpCUS1_gRNA_KI_BsaI_R | aaacAGATCTCGTCAGTAAGTGTC |
| SK0611 | PpCUS2_gRNA_KI_BsaI_F | ccatGGGACTTACTGACAAGATCG |
| SK0612 | PpCUS2_gRNA_KI_BsaI_R | aaacCGATCTTGTCAGTAAGTCCC |
| **Genotyping of CRISPR KO lines (PCR & Sanger sequencing)** | | |
| SK0448 | PpCUS1_ontarget2_F | TTCGGAAACCCTTCAGCAGG |
| SK0449 | PpCUS1_ontarget2_R | CGGAACCAAATGCACACTGC |
| SK0452 | PpCUS2_ontarget2_F | ACTTGCGTTGCGGTATCGTA |
| SK0453 | PpCUS2_ontarget2_R | AGCAGCAAACGGGACTTACT |
| **Genotyping of KI lines (PCR & Sanger sequencing)** | | |
| EL0197 | 5'UTR_CUS1_mScarlet3_F | TGGGTACAAGGAGCCAGTTA |
| EL0198 | 5'UTR_mScarlet3_R | CGCCGTCCTCGAAGTTCA |
| EL0199 | 5'UTR_CUS2_mScarlet3_F | CGCGAAGGACTGTGGGAAG |
| EL0202 | CUS1_CDS_3'UTR_R | ACTGTTGCTCACATGAAACTCTG |
| EL0208 | CUS1_CDS_3'UTR_F2 | CCCAGATCTCGTCAATGATTATCT |
| EL0210 | CUS2_CDS_3'UTR_R2 | TCCTTTGTCGCATTTGTGTTTTTG |
| EL0212 | CUS2_CDS_3'UTR_F3 | CCCCGATCTTGTCAATGATTACC |
| EL0035 | PTA2-3'UTR_R1 | TCTAGTTGGAGTCAGTTTCCCA |
| FB0159 | PTA2_5’UTR_F3 | AACTGTACGCAGTTCCGAGC |
| FB0165 | mNeonGreen-592-R | AACCAGCCGATGTACGTGTT |
| FB0167 | PTA2_5’UTR_F4 | GTTCCGAGCTGGTGAGTGAA |
| FB0168 | pCUS2_PTA2_R2 | TTACAGGTGAAGGACGAGCC |
| HR0627 | CYP73A49_sqF1 | TAATGCAGCGGTGTCGAGTT |
| HR0628 | CYP73A49_sqR1 | GCCGCGACGTTAATGTTCTC |
| HR0629 | CYP73A48_sqF1 | CGTGCAAGGATTATGCGTGG |
| HR0630 | CYP73A48_sqR1 | TCAGGCTCGCTCACCAAATT |

**Table S2. List of plasmids used in the study.**

| ID | Name | Purpose | Reference |
|---|---|---|---|
| **CRISPR** | | | |
| pHR0648 | pACT-Cas9 | SpCas9 expression | (Lopez-Obando et al. 2016) |
| pSK0024 | pENTR-U6P-sgRNA-CUS1_KO | sgRNA expression for CRISPR-assisted gene knock-out | This study |
| pSK0025 | pENTR-U6P-sgRNA-CUS2_KO | sgRNA expression for CRISPR-assisted gene knock-out | This study |
| pSK0249 | pENTR-U6P-sgRNA-PTA2_KI | sgRNA expression for CRISPR-assisted gene knock-in | This study |
| pSK0319 | pENTR-U6P-sgRNA-CUS1_KI | sgRNA expression for CRISPR-assisted gene knock-in | This study |
| pSK0320 | pENTR-U6P-sgRNA-CUS2_KI | sgRNA expression for CRISPR-assisted gene knock-in | This study |
| **Gene knock-in (KI)** | | | |
| pFB0003 | pGEM-T:CUS1pro:GUS-mNG-NLS | KI donor DNA for promoter-reporter strategy | This study |
| pFB0014 | pGEM-T:CUS2pro:GUS-mNG-NLS | KI donor DNA for promoter-reporter strategy | This study |
| pHR1064 | pGEM-T:SP-FLAG-mSC3-PpCUS1 | KI donor DNA for *cus1/cus2* mutant complementation | This study |
| pHR1066 | pGEM-T:SP-FLAG-mSC3-PpCUS2 | KI donor DNA for *cus1/cus2* mutant complementation | This study |
| ***P. patens* transformant selection** | | | |
| pHR0340 | pRT101:NPTII | G418-based selection of *P. patens* transformants | (Girke et al. 1998) |

**Table S3. List of quantification masses and molar response factors used for quantitative analysis of cuticular monomers by GC-TOFMS.** IS, internal standard; TMS, Trimethylsilyl group.

| **Analyte** | **Quantification mass** (±500ppm) | **Molar response factor** (relative to IS) |
|---|---:|---:|
| ribitol, 5TMS (IS for glycerol) | 73.08 | 1.00 |
| glycerol, 3TMS | 73.08 | 1.05 |
| C17:0, 1TMS (IS for methyl esters) | 73.06 | 1.00 |
| *t*-cinnamate, methyl ester | 131.06 | 2.11 |
| *p*-coumarate, 1TMS, methyl ester | 73.06 | 3.44 |
| caffeate, 2TMS, methyl ester | 219.05 | 0.84 |
| ferulate, 1TMS, methyl ester | 250.08 | 1.23 |
| sinapate, 1TMS, methyl ester | 280.09 | 4.79 |
| 16-OH C16:0, 1TMS, methyl ester | 75.05 | 2.13 |
| 10,16-diOH C16:0, 2TMS, methyl ester | 73.06 | 1.52 |
| 1,9,18-triOH C18:0, 3TMS | 73.06 | 1.52 |
| C14:0, methyl ester | 74.05 | 0.58 |
| C16:0, methyl ester | 74.05 | 0.61 |
| C18:0, methyl ester | 74.05 | 0.61 |
| C20:0, methyl ester | 74.05 | 0.62 |
| C22:0, methyl ester | 74.05 | 0.62 |
| C24:0, methyl ester | 74.05 | 0.64 |
| C18:1, methyl ester | 55.09 | 1.16 |
| C18:2, methyl ester | 67.08 | 0.25 |
| C18:3, methyl ester | 79.07 | 0.76 |
| C20:4, methyl ester | 79.07 | 0.62 |
